# MeTAL enables multiparametric risk prediction for human *KCNH2* variants

**DOI:** 10.64898/2026.04.29.721778

**Authors:** Barbara Ribeiro de Oliveira, Elouan Voisin, Maxence Marbouty, Achille Gregoire, Malak Alameh, Jérôme Montnach, Aurélie Thollet, Flavien Charpentier, Isabelle Denjoy, Vincent Probst, Gildas Loussouarn, Isabelle Baró, Michel De Waard, Rupamanjari Majumder

## Abstract

**Background:** Clinical interpretation of missense variants in the hERG potassium channel encoded by the *KCNH2* gene remains a major challenge in inherited arrhythmia syndromes. Functional studies often rely on a minimal set of channel properties, mainly current amplitude measurements, which do not capture the multidimensional nature of channel gating and its impact on ventricular repolarization. We developed a multiscale computational framework to quantitatively link multiparametric channel dysfunction to ECG phenotypes.

**Methods:** We generated multiparametric electrophysiological profiles for *KCNH2* variants using high-throughput patch-clamp, quantifying nine biophysical properties including conductance, voltage dependence, and gating kinetics. These parameters were incorporated into a modified formulation of I_Kr_ embedded in a human ventricular electrophysiology model. The resulting framework, termed MeTAL (Multiscale-enriched Transformation and Analysis for Long-QT), produces physiologically calibrated pseudo-ECGs enabling quantitative evaluation of QT dynamics. We systematically analyzed the contribution of individual and combined gating parameters and applied the model to 41 *KCNH2* variants across ACMG classes, comparing simulated QTc values with clinical data.

**Results:** Multiparametric profiling revealed complex functional signatures in most variants, with concurrent gain- and loss-of-function effects affecting distinct gating processes. Simulations demonstrated that ventricular repolarization, though strongly determined by current amplitude, is substantially influenced by inactivation-related parameters, particularly the slope and voltage dependence of inactivation. Interaction analyses showed nonlinear relationships between gating parameters, explaining why variants with similar current density can produce divergent QT phenotypes. In heterozygous simulations, MeTAL reproduced clinically observed QTc distributions across variant classes and accurately predicted the repolarization regime (normal, long-QT, or short-QT) in most cases.

**Conclusions:** Multiparametric integration of ion-channel function within a multiscale electrophysiological model enables mechanistic prediction of QT behavior beyond conductance-based metrics. This approach provides a scalable framework for interpretation of *KCNH2* variants and solves the issue of risk stratification in inherited arrhythmia syndromes while offering new opportunities for variant-specific and pharmacological modeling of repolarization.

## Introduction

Short- and long-QT syndromes (SQTS and LQTS) are life-threatening hereditary or acquired heart rhythm disorders associated with syncope, ventricular arrhythmias, and sudden cardiac death. Missense mutations in *KCNH2*, the gene encoding the ether-à-go-go hERG potassium channel, represent a major genetic substrate of inherited LQT2 and are increasingly identified through genetic testing. ^1^ However, interpretation of these variants remains a major clinical challenge. Of the 3,909 *KCNH2* variants currently catalogued in ClinVar (*https://www.ncbi.nlm.nih.gov/clinvar/* on the 29^th^ of April, 2026), 1,870 are missense variants, among which approximately 80% are classified as variants of uncertain significance (VUS), limiting their utility for diagnosis, risk stratification, and therapeutic decision-making. Several high-throughput functional initiatives have therefore been launched to improve pathogenicity assessment.^2,3^

In current practice, functional interpretation of hERG variants frequently relies on measurements of current density, for example, through maximal conductance (G_max_) or peak tail current, which can provide an integrated readout of trafficking and major gating defects.^3,4,5^ While this approach is effective for identifying severe loss-of-function variants, it necessarily compresses the multidimensional behavior of hERG gating into a limited set of quantitative metrics. The contribution of hERG channels to ventricular repolarization depends not only on current amplitude, but also on the voltage dependence and kinetics of activation, inactivation, and deactivation, each of which has the power to independently reshape the time course of the hERG current (*I*_*Kr*_) and thereby influence repolarization. Prior computational and experimental studies have highlighted the importance of these gating processes.^5,6^ However, because of the multiscale nature of the problem (ion channel → cell → tissue → organ → patient), the relative quantitative contributions to clinical pathogenicity remain incompletely defined. Moreover, the state-of-the-art mechanistic frameworks that do link multiparametric channel dysfunction to individual electrophysiological phenotypes are computationally suboptimal for treating a large set of *KCNH2* variants. This gap is particularly relevant for variants that produce mixed or subtle functional alterations that may not be fully captured by integrated current measurements alone.

In this context, mathematical modeling provides a powerful framework to bridge ion channel dysfunction and cardiac electrophysiology across scales. The two principal classes of mathematical models used to describe ion channel behavior are the Hodgkin-Huxley (HH) formulation and Markov state models.^7,8^ HH-type models are widely adopted because of their relative simplicity and ease of parameterization. However, their structure inherently couples voltage dependence and kinetics, thereby limiting flexibility in representing disease-associated gating perturbations. In contrast, Markov models offer greater mechanistic detail but are often difficult to parameterize directly from experimental data. Importantly, many existing hERG formulations rely on simplifying assumptions – such as instantaneous inactivation – that obscure the contribution of inactivation dynamics, a property frequently altered in disease-associated variants.^9,10^ For variant interpretation, an optimal framework should seamlessly integrate high-throughput functional measurements with patient-level classification, while accurately capturing the full spectrum of hERG channel biophysical properties. These limitations underscore the need to develop advanced modeling strategies capable of systematically incorporating multiparametric channel dysfunction into physiologically meaningful predictions.

To address these challenges, we introduce MeTAL (Multiscale-enriched Transformation and Analysis for Long-QT), a mechanistically grounded framework that integrates a nine-parameter formulation of *I*_*Kr*_ into an established multiscale model of human ventricular electrophysiology.^8^ Built on an adapted HH structure that uncouples channel kinetics from voltage dependence, the model interfaces directly with high-throughput generated functional data and preserves all the key features of hERG gating. ^11-15^ The framework is coupled to a standard pseudo-ECG generation pipeline,^16^ followed by a nonlinear transformation module that temporally rescales and realigns the simulated signals to produce physiologically plausible ECG traces corresponding to leads V5 and V6 of the standard 12-lead ECG, enabling quantitative assessment of variant effects on QT dynamics. Using this framework, we first quantified the relative contributions of individual hERG biophysical parameters to ventricular repolarization. We then examined how simultaneous changes in two parameters influence repolarization, capturing more complex variant effects. Finally, we evaluated model performance across 41 *KCNH2* variants spanning ACMG classes 1-2 (benign/likely benign, n=21) and classes 4-5 (likely pathogenic/pathogenic, n=20), and compared simulated and clinical QT intervals in patient cohorts carrying representative variants – 2 class 1 variants (4 patients), 5 class 2 variants (11 patients), 8 class 4 variants (10 patients), and 12 class 5 variants (33 patients). Our findings identify inactivation-related parameters, alongside hERG current amplitude, as dominant determinants of ventricular repolarization. Multiparametric functional profiling improves mechanistic interpretation and prediction of QT risk beyond conductance-based metrics alone. By integrating experimental ion-channel measurements with multiscale electrophysiological simulations and morphology-aware ECG transformation, MeTAL connects channel-level dysfunction to clinically interpretable ECG phenotypes. This approach enables flexible ECG representations and provides a scalable framework for quantitative interpretation of *KCNH2* variants in clinical genomics and arrhythmia risk assessment.

## Results

### Clinical and multiparametric heterogeneity of KCNH2 variants

Marked interpatient variability in ECG recordings, particularly in QTc duration and stability, is a well-recognized feature of *KCNH2*-mediated long-QT syndrome.^17^ Clinical V5 ECGs are shown for four LQT2 patients with different hERG missense variants (**Figure 1A**). Phenotypic variation in QTc among LQT2 patients reflects, among many other factors, the multidimensional contributions of hERG channel expression and gating to ventricular repolarization. To capture this complexity, we conducted standardized high-throughput electrophysiological profiling of representative KCNH2 variants, quantifying nine key biophysical parameters spanning current amplitude, voltage dependence – half-maximal activation and inactivation potentials (V_1/2_) and slope factors (k) of steady-state activation and inactivation – and gating kinetics of activation, deactivation, inactivation, and recovery from inactivation.^12^ These parameters define each variant multidimensional functional profile, which we visualize as radar plots for eight biophysical parameters and individual plots for current amplitude (**Figure 1B**). In the radar plot representation, each axis denotes the deviation of a parameter from the wild-type value – expressed as absolute differences for V_1/2_ and as ratios for the remaining parameters. For details on the interpretation of radar plots, see **Supplementary Methods**.

**Figure 1.**
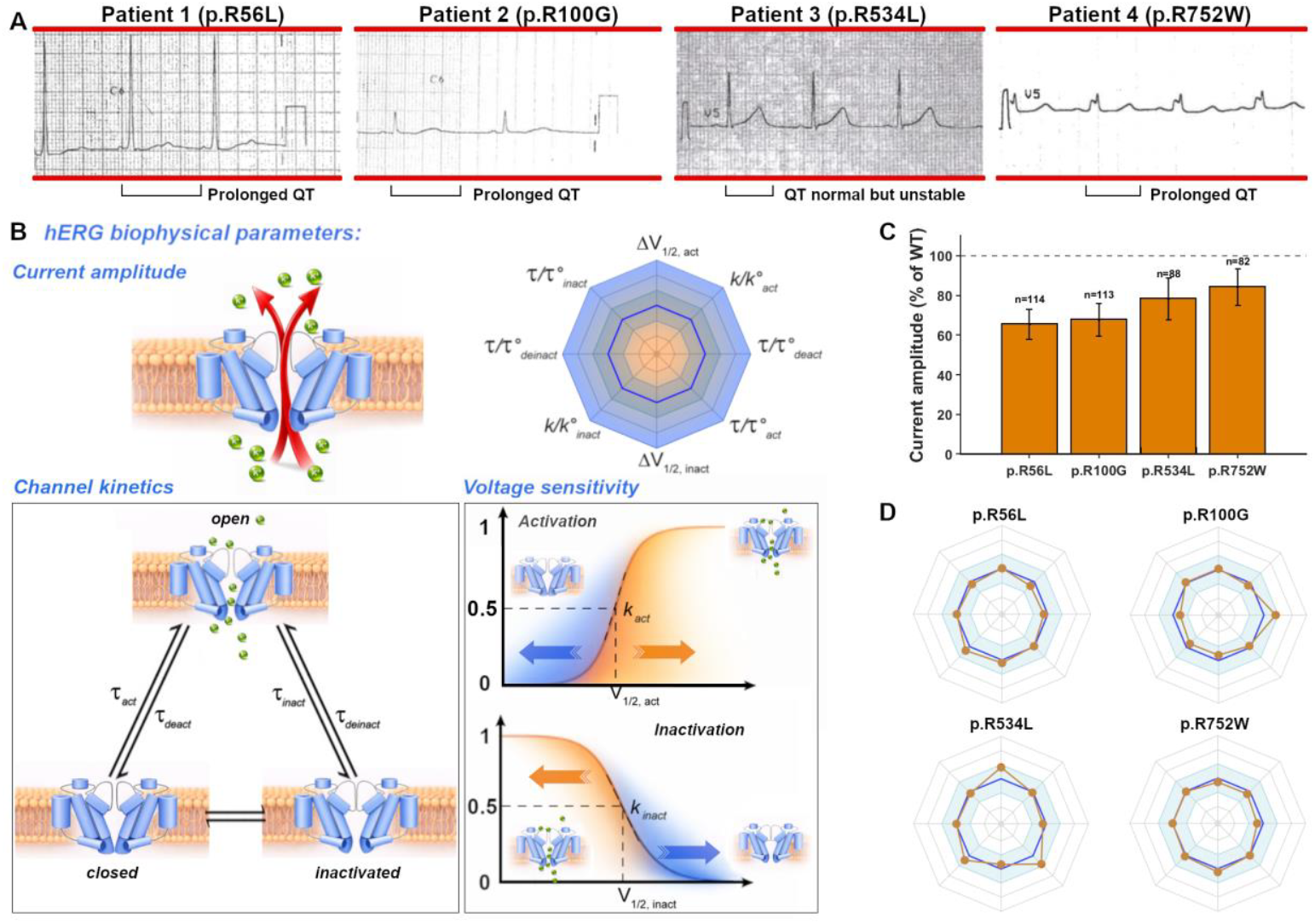
Interpatient variability and multiparametric functional remodeling of *KCNH2* variants. **A**, Representative clinical ECG recordings illustrating the wide range of QT phenotypes observed among patients carrying *KCNH2* variants, including markedly prolonged and near-normal QT intervals. **B**, Schematic guide for interpreting the nine-parameter radar (octagonal) representation of hERG function. Each axis corresponds to one biophysical parameter, including voltage dependence (V_1/2_), slope (k), and gating kinetics (*τ*) expressed as time constants, except for deactivation, expressed as time to decrease to 50% of the maximum value. The wild-type reference is shown with a dark blue line. Color coding (orange, loss of function; blue, gain of function) and radial displacement relative to the wild-type octagon indicate the direction and magnitude of variant-induced changes (see Supplementary material for more details). Current amplitude is not represented in the radar plot since gating properties are unknown for variants with complete amplitude loss of function. **C**, Bar plot showing the percentage of current amplitude retained in the corresponding hERG variants relative to wild-type (WT). **D**, Radar plots showing heterozygous multiparametric functional profiles of representative *KCNH2* variants as brown filled circles normalized to WT (blue octagon zone). Outward deflections indicate gain of function, whereas inward deflections indicate loss of function.

Multiparametric profiling of the *KCNH2* variants shown for the four patients revealed that most produced complex functional fingerprints (**Figure 1C, D**). Individual variants showed manifest overall reductions in hERG current amplitude in heterozygous conditions, along with gain- and loss-of-function effects across distinct gating properties (**Figure 1D**). Notably, alterations were distributed across activation and inactivation processes, with several variants showing substantial perturbations in parameters not routinely captured by conductance-centric assessments. For instance, the variant p.R534L displayed a modest reduction in current amplitude but was accompanied by marked shifts in inactivation-related parameters, whereas p.R56L combined smaller alterations occurring across multiple parameters.

These observations raise two key questions: (i) what is the relative contribution of individual biophysical parameters to pathogenicity, and (ii) how should their combined effects be interpreted when parameters exert opposing gain- and loss-of-function influences on overall risk? Addressing these challenges requires integrative frameworks capable of systematically linking channel-level perturbations to repolarization of phenotypes. These questions motivated us to develop **MeTAL** (Multiscale-enriched Transformation and Analysis for Long-QT), a simulation platform for quantitative analysis of *KCNH2* variant effects.

### Development of a novel mathematical formulation of the hERG channel function to be integrated in MeTAL

In MeTAL we aimed to incorporate an improved mathematical formulation of hERG channel electrophysiology within a multiscale adaptation of the O’Hara-Rudy (ORd) ventricular model^9^. The revised mathematical formulation of hERG channel gating is fully described in **Methods** and **Supplementary Methods** and briefly summarized below. The total hERG current *I*_*Kr*_ was reformulated according to Eq.1:

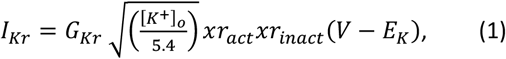

Within the gating formulation, we retained the original equations for the voltage dependence of steady state activation (*xr*_*act*,∞_) from the ORd model (**Figure 2A**), while replacing the kinetics (*τ*_*act*_) with Eqs.2-5, for the fast (*τ*_*xrf*_) and slow (*τ*_*xrs*_) components (**Figure 2B**).

**Figure 2.**
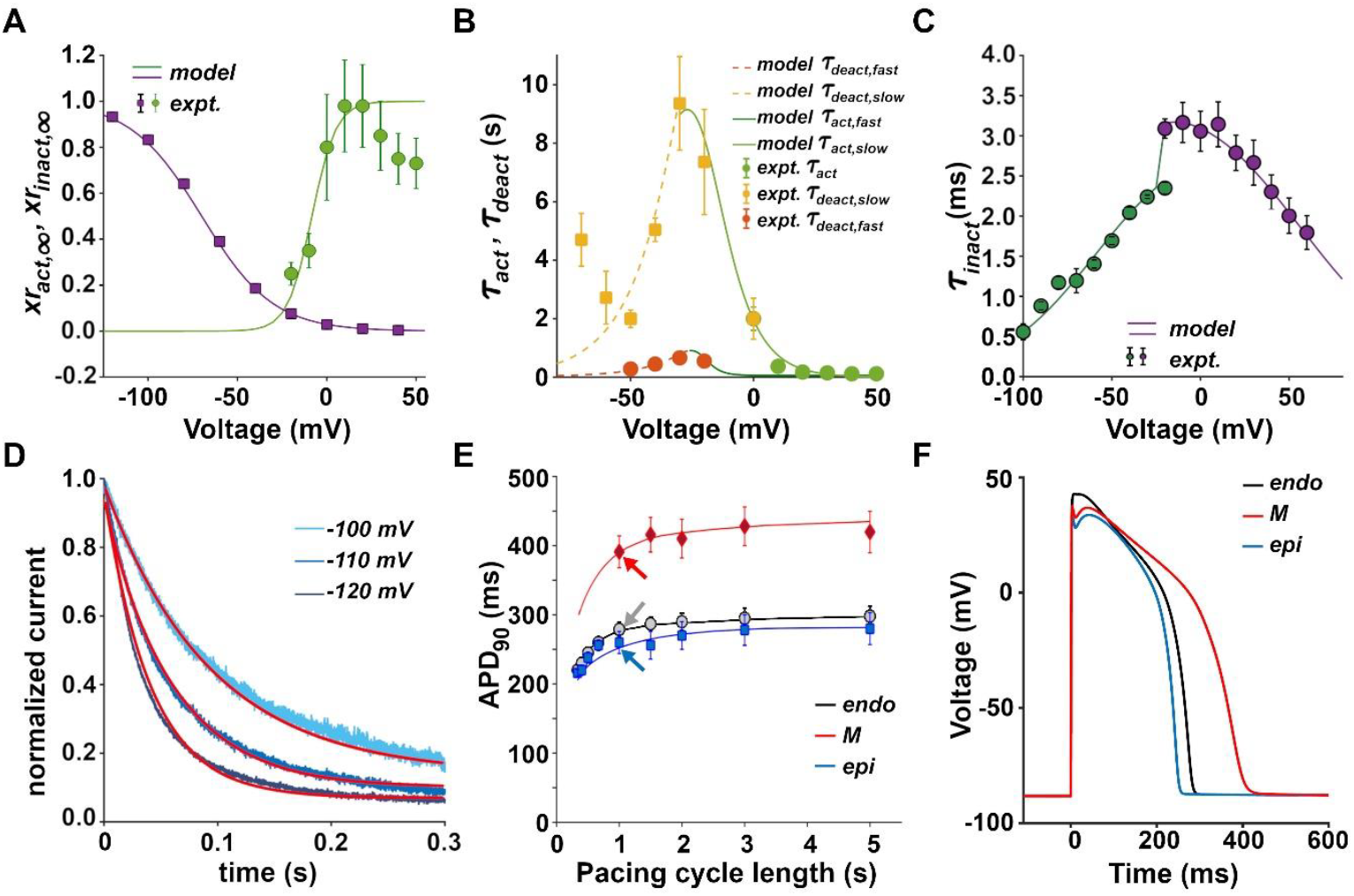
Validation of the hERG formulation and resulting ventricular action potentials. Biophysical properties of the wild-type hERG channel and resulting action potentials as implemented in MeTAL. **A**, Steady-state voltage dependence of hERG activation and inactivation. Experimental measurements (symbols^9,18^) and simulated values (lines) are shown for activation and inactivation gating. **B**, Voltage dependence of activation and deactivation time constants. Symbols denote experimental measurements^9^ and lines denote simulated values. **C**, Voltage dependence of kinetic of inactivation (purple) and recovery from inactivation (green). Symbols represent experimental measurements^19^ and lines represent simulated values. **D**, Normalized deactivation currents measured at 22°C at -120, -110, and -100 mV (blue lines) and fitted with exponentials yielding time constants (*τ*) of 40 ms, 60 ms, and 94 ms, respectively.^13^ Simulated values (red lines) reproduce experimentally observed kinetics after temperature correction. **E**, Restitution of action potential duration at 90% repolarization (APD_90_) as a function of pacing cycle length in epicardial, midmyocardial, and endocardial cells. Symbols indicate experimental measurements^9,20^ and lines indicate simulated results (arrows denote APD_90_ at a pacing cycle length of 1000 ms). **F**, Representative simulated action potentials in subendocardial (black), midmyocardial (red), and subepicardial (blue) cells paced at a cycle length of 1000 ms.

For *V* ≥ −30*mV*

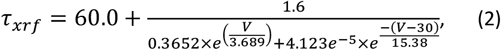

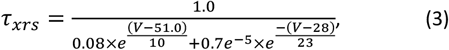

Otherwise,

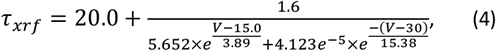

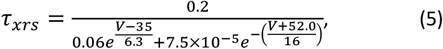

The net kinetic of activation *τ*_*act*_ is a weighted sum of these two components, as proposed within the framework of the ORd model. We also reformulated the inactivation gating by introducing a voltage-dependent steady state (*xr*_*inact*,∞_) according to Eq. 6 (**Figure 2A**):

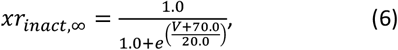

And included a kinetic of inactivation, given by Eqs.7-11 (**Figure 2C**).

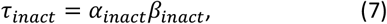

For *V* ≥ −20.0*mV*

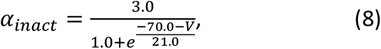

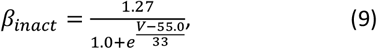

Otherwise,

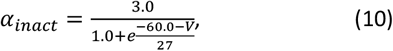

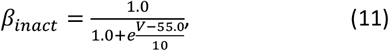

Note that, in experiments, the *τ*_*xrs*_ and *τ*_*xrf*_ cannot be reliably uncoupled; instead, the measured signal reflects *τ*_*act*_. Therefore, to link the model to experimental data, and to uncouple the kinetics of activation from the kinetics of deactivation, we used the following rule: *τ*_*xrs*_ = *τ*_*xrs*_ × Γ_1_, where Γ_1_ = Γ_*deact*_ for *V* < −30 *mV* and Γ_*act*_, otherwise. Similarly, *τ*_*xrf*_ = *τ*_*xrf*_ × Γ_2_, where Γ_2_ = Γ_*deact*_ for *V* < −26 *mV*, and Γ_*act*_, otherwise.

Introducing this new hERG formulation required additional adjustments to the maximal conductance of several other cardiac ion channels to preserve ventricular repolarization, particularly restitution properties (for details, please refer to **Supplementary Methods**).

Thanks to this new mathematical formulation, it is now possible to independently and simultaneously control the nine experimentally measurable hERG parameters (**Figure 1B**). Next, we validated the performance of the mathematical formulation by quantitative comparison with experimental data. Simulated activation and deactivation kinetics matched previously reported voltage-dependent time constants measured at physiological temperature (**Figure 2D**). Simulated APD_90_ restitution curves reproduced experimentally reported relationships between pacing cycle length and action potential duration across epicardial, midmyocardial, and endocardial cell types (**Figure 2E**). Finally, the mathematical formulation allowed the generation of stable action potentials with expected morphology at a pacing cycle length of 1000 ms (**Figure 2F**), including the characteristic absence of a notch in sub-endocardial cells.^20,21^ Together, these results confirm that the revised hERG mathematical formulation preserves physiologically realistic ventricular electrophysiology while enabling independent manipulation of experimentally measurable channel parameters, providing the foundation for quantitative simulation of variant-specific effects on ventricular repolarization. **Table S1** summarizes the general definitions, abbreviations, and wild-type reference values for all the parameters used in the mathematical formulation. **Table S2** lists the values of the parameters that were altered to generate ventricular action potentials. Now that a robust mathematical formulation of the hERG channel gating has been established based on the entire set of biophysical properties, we next proceeded with pseudo-ECG generation.

### Pseudo-ECG generation and recalibration

To derive clinically interpretable signals from the cellular electrophysiological simulations described above, pseudo-ECGs were first generated using the classical Gima-Rudy approach^16^ with the ORd single-cell models. Mathematical reformulation of hERG channel properties improved the stability of the ventricular cardiomyocyte model, rendering it more computationally efficient and robust than the original ORd model (**Figure S1**). However, comparison with clinical biomarker values^22^ for wild-type hERG channel indicates that this approach does not eliminate the systematic distortions in baseline pseudo-ECG morphology, consistent with those observed in current state-of-the-art models of clinically coherent pseudo-ECG generation. Consequently, simulated pseudo-ECGs exhibit prolonged ST segments and shortened T-wave durations, and the resulting in QT intervals clustered near the lower bound of the physiological range (**Supplementary Methods**). These limitations reduce the physiological interpretability of simulated signals and may compromise their ability to accurately recapitulate patient-specific ECG phenotypes.

This issue likely reflects the highly reduced, one-dimensional nature of the Gima-Rudy formulation, which may be overly simplified and therefore susceptible to numerical artefacts, including the systematic distortions in biomarker values observed here. We therefore hypothesized that applying a nonlinear recentering transformation to the pseudo-ECG, without modifying the underlying electrophysiological model which had already been optimized, would be sufficient to restore agreement with clinical ECG characteristics. Accordingly, prior to interpreting the clinical impact of hERG variants, recalibration of the pseudo-ECG is required to accurately represent the wild-type condition. To this end, we applied a segment-wise transformation to the Gima–Rudy pseudo-ECG, rescaling and recentering individual intervals while preserving waveform morphology (**Figure 3, S2)**. Recalibration was performed using 2,500 transmural ventricular strand configurations (**Figure S2A**) incorporating baseline hERG parameters (**Figure 2A-D**) and intercellular variability (**Figures 3A**). Pseudo-ECGs were computed, and fiducial Q, R, S, and T points were identified to extract ECG biomarkers, including QRS duration, ST segment duration, T-wave duration, and QT interval (**Figure 3B**). Signals with abnormal morphologies (e.g, missing or repeated fiducial points) were excluded (**Figure 3C**). From the remaining population, median values for each biomarker were computed (**Figure S2D, E**). Scaling coefficients were derived as the ratio of sampled physiological biomarker values to their medians and used to linearly rescale individual pseudo-ECG segments, ensuring physiological consistency (**Figure 3D**). The rescaled segments were subsequently concatenated using spline interpolation to generate the recalibrated ECG (**Figure 3D**, lower panel).

**Figure 3.**
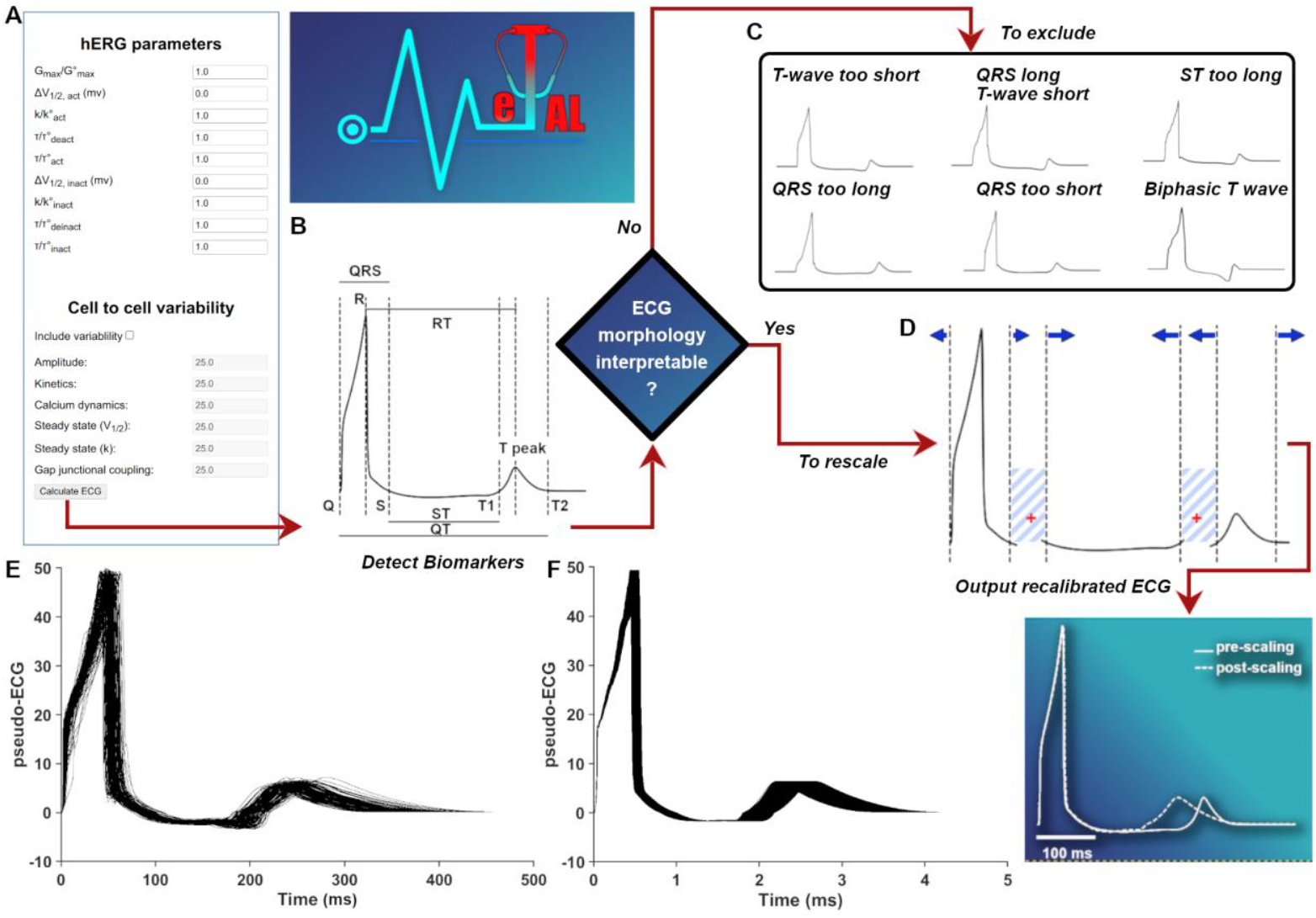
Generation and calibration of pseudo-ECGs for physiologically consistent biomarker extraction. **A**, MeTAL workflow with hERG parameter perturbations as inputs (left, software interface view). **B**, Pseudo-ECG with fiducial Q, R, S, T_1_ and T_2_ points for automated extraction of QRS, ST segment, T-wave (T_2_-T_1_), and QT durations. **C-D**, Segment-wise calibration. Scaling coefficients derived from 2,500 transmural strand simulations were used to rescale and recenter ECG intervals while preserving waveform morphology. Nonphysiological configurations were excluded (**C**), and segments recombined using spline interpolation (blue stripes, **D**) to output a recalibrated pseudo-ECG. **E**, Representative pseudo-ECGs illustrating the effect of intercellular variability on ECG morphology under normal conditions. Recalibration was performed using scaling coefficients derived from physiological mean values of each biomarker. **F**, Representative set of 1,000 recalibrated pseudo-ECGs obtained from a single simulated transmural strand configuration using 1,000 distinct combinations of permissible scaling coefficients, generated by uniformly sampling biomarker values within their physiological ranges.

Two complementary sources of diversity were introduced during recalibration. First, intercellular variability was incorporated to generate intrinsic differences in pseudo-ECG morphology (**Figure 3E**; in this case, scaling coefficients were derived from physiological mean biomarker values). Second, diversity was further expanded by uniformly sampling a large number of biomarker values within their physiological ranges to compute scaling coefficients. **Figure 3F** shows 1,000 representative recalibrated pseudo-ECGs from a single transmural strand configuration, obtained using 1,000 distinct combinations of permissible scaling coefficients. This procedure yielded pseudo-ECGs that closely reproduce the timing and morphology of V5/V6 leads of a standard 12-lead ECG in a normal individual.

The next step was to assemble an integrated pipeline linking hERG channel electrophysiology to pseudo-ECG generation and clinically coherent recalibration within a unified, user-friendly, academic open-source computational platform. This platform, termed MeTAL, establishes a standardized baseline representation of ventricular repolarization for the wild-type *I*_*Kr*_ current and enables quantitative assessment of variant-induced perturbations in ECG biomarkers within physiologically realistic ranges. The full assembly of the MeTAL pipeline is described in **Supplementary Methods** and in **Figure S2**. By enabling controlled multiparametric perturbations of hERG gating and direct evaluation of their downstream ECG consequences, MeTAL provides a structured framework for linking variant-specific channel dysfunction to clinically interpretable ECG phenotypes.

### Evaluation of pathogenicity of single hERG parameter variations using MeTAL

To evaluate the pathogenic potential of individual hERG perturbations, we systematically varied each of the nine experimentally measurable channel parameters while keeping all others fixed. For each perturbation, pseudo-ECG signals were generated and analyzed using MeTAL to quantify effects on QT duration and ventricular repolarization (summarized in **Figure 4A**; see Methods).

**Figure 4.**
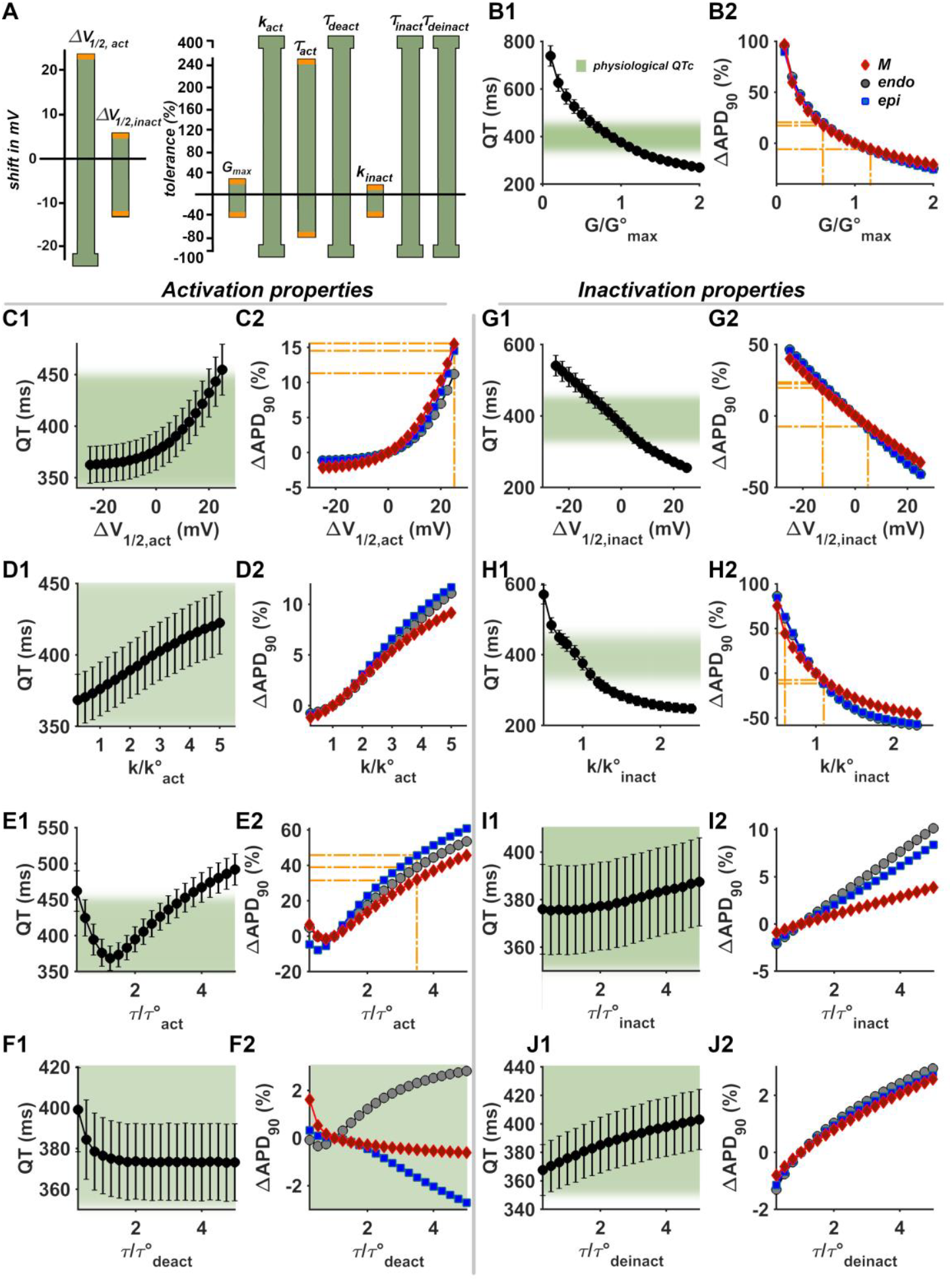
Contribution of individual hERG parameters to QT interval and APD_90_ as assessed using the MeTAL framework. **A**, Physiological tolerance ranges for each hERG parameter derived from single-parameter perturbations. Bars indicate the range of values over which QT intervals remain within physiological limits; Hammer heads denote ranges extending beyond the tested parameter space. Orange boundaries indicate approximate thresholds for transition to abnormal QT behavior. **B**, Effect of changes in maximal conductance (G/G°_max_) on QT interval (mean±SD; **B1**) and APD_90_ (**B2**). Shaded regions indicate physiological QT ranges. Orange lines denote APD_90_ values at the threshold of QT abnormality for each cell type. **C-F**, Effects of activation and deactivation parameters (V_1/2,act_, k_act_, *τ*_act_, *τ*_deact_) on QT interval and APD_90_. **G-J**, Effects of inactivation and recovery parameters (V_1/2,inact_, *k*_inact_, *τ*_inact_, *τ*_deinact_) on QT interval and APD_90_. Endo, subendocardial; M, midmyocardial; epi, subepicardial cells.

Variation of maximal conductance (G_max_) produced the expected effects on repolarization. QT prolongation occurred for G/G°_max_ ≤ 0.6 (*i*.*e*. a decrease of at least 40% in **Figure 4A**), whereas QT shortening was observed for G/G°_max_ ≥ 1.2 (**Figure 4B1**). These changes were consistent across cell types and corresponded to increase (∼17-19%) or decrease (∼6%) in APD_90_ (**Figure 4B2**, and **Figure S3A** for details at 3 different G/G°_max_ values).

Modulation of activation properties produced more complex effects. Positive shifts in V_1/2,act_ beyond +25 mV resulted in QT prolongation, whereas negative shifts within the explored range did not induce pathological QT changes (**Figure 4C**; **Figure S3B**). Variations in *k*_act_ had minimal impact, with APD_90_ changes remaining modest (<15%; **Figure 4D**; **Figure S3C**). In contrast, slowing activation kinetics (*τ*_act_) produced QT prolongation, corresponding to 30-45% increases in APD_90_ and exhibiting regional heterogeneity across ventricular cell types (**Figure 4E**; **Figure S3D**).

However, inactivation properties exerted the strongest influence on repolarization. Negative shifts in V_1/2,inact_ beyond -12.5 mV led to QT prolongation, whereas positive shifts exceeding +5 mV produced QT shortening (**Figure 4G**), accompanied by APD_90_ changes exceeding ∼20% increase or ∼8% reduction (**Figure S4A**). Among all parameters, the slope of inactivation (*k*_inact_) emerged as the most effective determinant of QT behavior. Relatively small perturbations were sufficient to induce pathological QT changes, due to marked alterations in APD_90_ (**Figure 4H**; **Figure S4B**). In contrast, variations in the kinetics of inactivation and recovery from inactivation had comparatively little effects (**Figure 4I, J**; **Figure S4C, D**).

Overall, hERG parameters exhibited marked heterogeneity in their tolerance to perturbation. While several parameters tolerated large deviations without producing abnormal QT intervals, others – including G_max_, V_1/2,inact_, *τ*_act_, and particularly *k*_inact_ – showed narrow physiological tolerance ranges. Notably, *k*_inact_ represents a relatively understudied aspect of hERG gating yet emerged here as a dominant determinant of repolarization. Importantly, the relationship between APD_90_ and QT varied across parameters, indicating that APD_90_ alone is insufficient to classify variant pathogenicity.

These findings provide a mechanistic basis for the multidimensional functional effects observed in *KCNH2* variants.

### Influence of hERG activation and inactivation parameters on QT sensitivity to G_max_

Because most *KCNH2* variants affect multiple biophysical properties, we next examined how combinations of hERG parameters interact to determine QT behavior. Using MeTAL, we systematically evaluated the joint effects of G_max_ and each gating parameter changes, including scenarios in which parameters changed in opposite directions (**Figure 5**; APD_90_ analysis in **Figure S5**).

**Figure 5.**
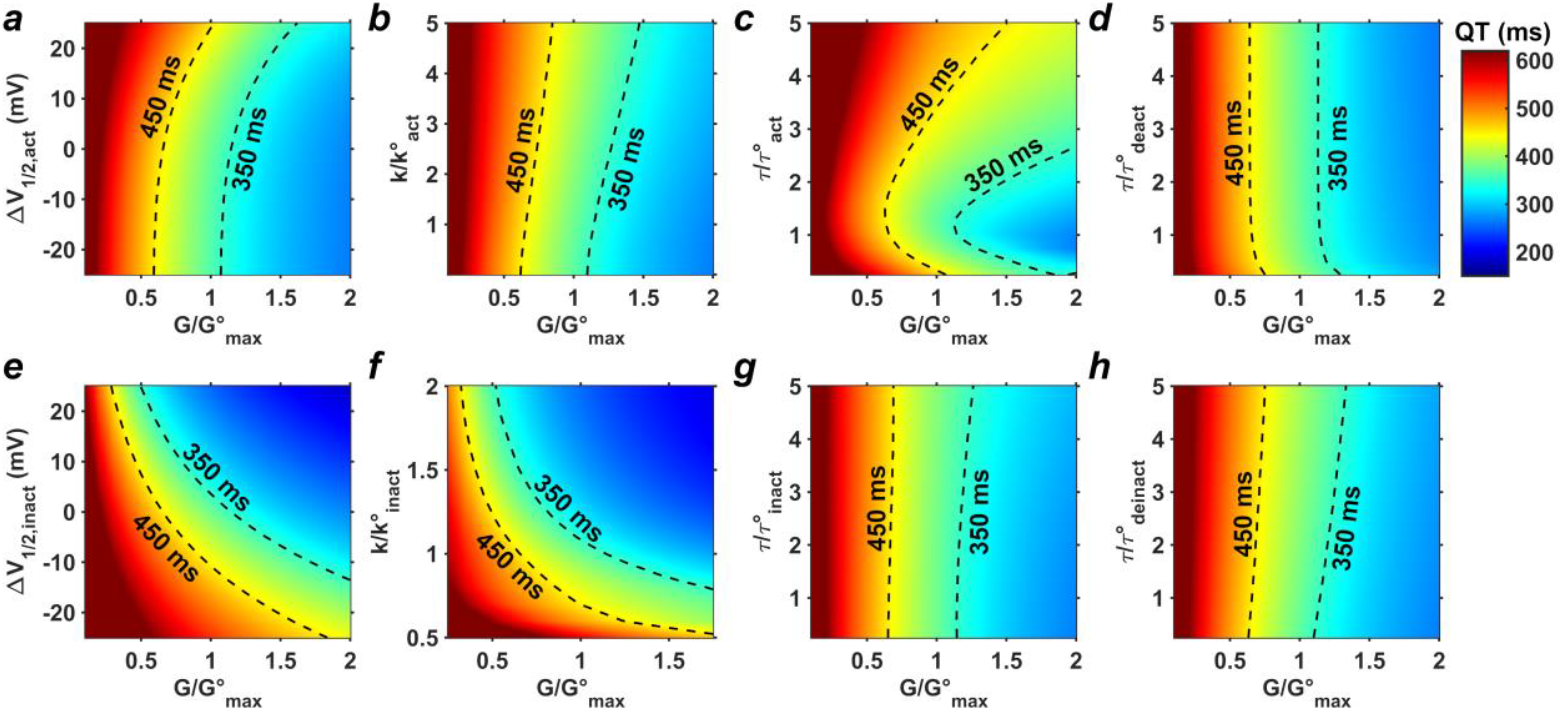
Pseudocolor maps of QT duration as a function of paired hERG biophysical parameters, including maximal conductance (G/G°_max_). Dashed lines delineate the boundaries of the physiological QT range. Color coding indicates QT phenotype, with blue representing QT shortening and red representing QT prolongation; color intensity reflects the magnitude of deviation from the physiological range.

When activation properties were varied in combination with G_max_, the resulting effects on QT duration were heterogeneous but generally remained dominated by G_max_. Positive shifts in V_1/2,act_ facilitated QT prolongation at less reduced G_max_ values only (**Figure 5A**), and variations in *k*_act_ had minimal impact (**Figure 5B**). For example, a +15-mV shift in V_1/2,act_ resulted in QT prolongation at G/G°_max_ ≈ 0.8. In contrast, slowing activation kinetics (*τ*_act_) markedly precipitated QT prolongation, even at normal G°_max_, due to to 30-45% APD_90_ increases that exhibit regional heterogeneity across ventricular cell types (**Figure 5C**; **Figure S5**). Deactivation kinetics had little influence on QT behavior across the explored range (**Figure 5D**).

In contrast, inactivation properties exerted a dominant influence over the effect of G_max_ on QT duration. Negative shifts in V_1/2,inact_ strongly promoted QT prolongation, such that even substantial increases in G_max_ failed to restore physiological QT values (**Figure 5E**). For instance, shifts beyond -20 mV resulted in prolonged QT despite increases in G_max_ of up to 1.5-fold. Conversely, positive shifts produced QT shortening even at normal conductance levels. Similarly, reductions in the slope of inactivation (*k*_inact_) markedly increased QT duration and could override the effects of increased G_max_, whereas relatively small increases in *k*_inact_ were sufficient to induce short QT (**Figure 5F**). These changes were accompanied by substantial alterations in APD_90_, consistent with the strong sensitivity of repolarization to inactivation dynamics (**Figure S5**). In contrast, variations in the kinetics of inactivation and recovery from inactivation had comparatively modest effects (**Figure 5G, H**; **Figure S5**).

These interaction patterns demonstrate that the width and shape of the physiological “safety zone” for QT duration depend critically on the underlying combination of gating parameters. In particular, inactivation parameters substantially narrow this zone, such that small perturbations can shift the system toward either long- or short-QT phenotypes. Notably, the concave shape of the safety boundary observed for certain parameter combinations – such as activation kinetics *versus* current amplitude – illustrates how QT prolongation can arise even in regions where increased G_max_ alone would predict shortening. These findings illustrate how similar degrees of hERG dysfunction can give rise to divergent QT outcomes depending on the underlying parameter combination and are consistent with our observation that MeTAL accurately reproduces QTc values measured in patients carrying *KCNH2* variants.

Overall, these results indicate that the impact of G^max^ on QT duration depends strongly on its interaction with other hERG parameters. Although G_max_ is the most studied determinant of channel function, it does not uniformly predict repolarization outcomes. Instead, activation kinetics and, more prominently, inactivation properties – particularly the slope of inactivation (*k*_inact_) – emerge as strong modulators of QT sensitivity. These findings underscore the importance of multiparametric functional assessment and demonstrate that reliance on a few parameters or APD_90_, is insufficient for accurate classification of variant pathogenicity.

### Use of MeTAL as a prescreen tool for variant assessment

Building on these insights, we next used MeTAL to investigate how multiparametric alterations reflecting those observed experimentally jointly influence QT duration. Variants in ion channels often affect multiple biophysical properties simultaneously, with some alterations contributing to pathogenicity while others may be functionally neutral.

Under heterozygous conditions, which reflect the clinical setting, the intrinsic effects of a variant may be obscured by the complexity of the patient-specific electrophysiological environment. As not all contributing factors can be captured in a computational model, isolating the direct impact of channel dysfunction remains challenging. To address this, we performed an initial prescreening of variants under homozygous conditions to assess their intrinsic pathogenic potential. We analyzed 41 *KCNH2* variants spanning ACMG classes 1 (n=5), 2 (n=16), 4 (n=8), and 5 (n=12). Initial multiparametric electrophysiological profiling of those variants in homozygous condition by high-throughput patch-clamp revealed substantial heterogeneity across variants, with concurrent gain- and loss-of-function effects distributed across the nine hERG biophysical parameters (**Figure 6A-B, S6**, and **Table S3**). Variants from classes 4-5 generally showed larger deviations from wild-type behavior, whereas variants from classes 1-2 exhibited often more modest alterations.

**Figure 6.**
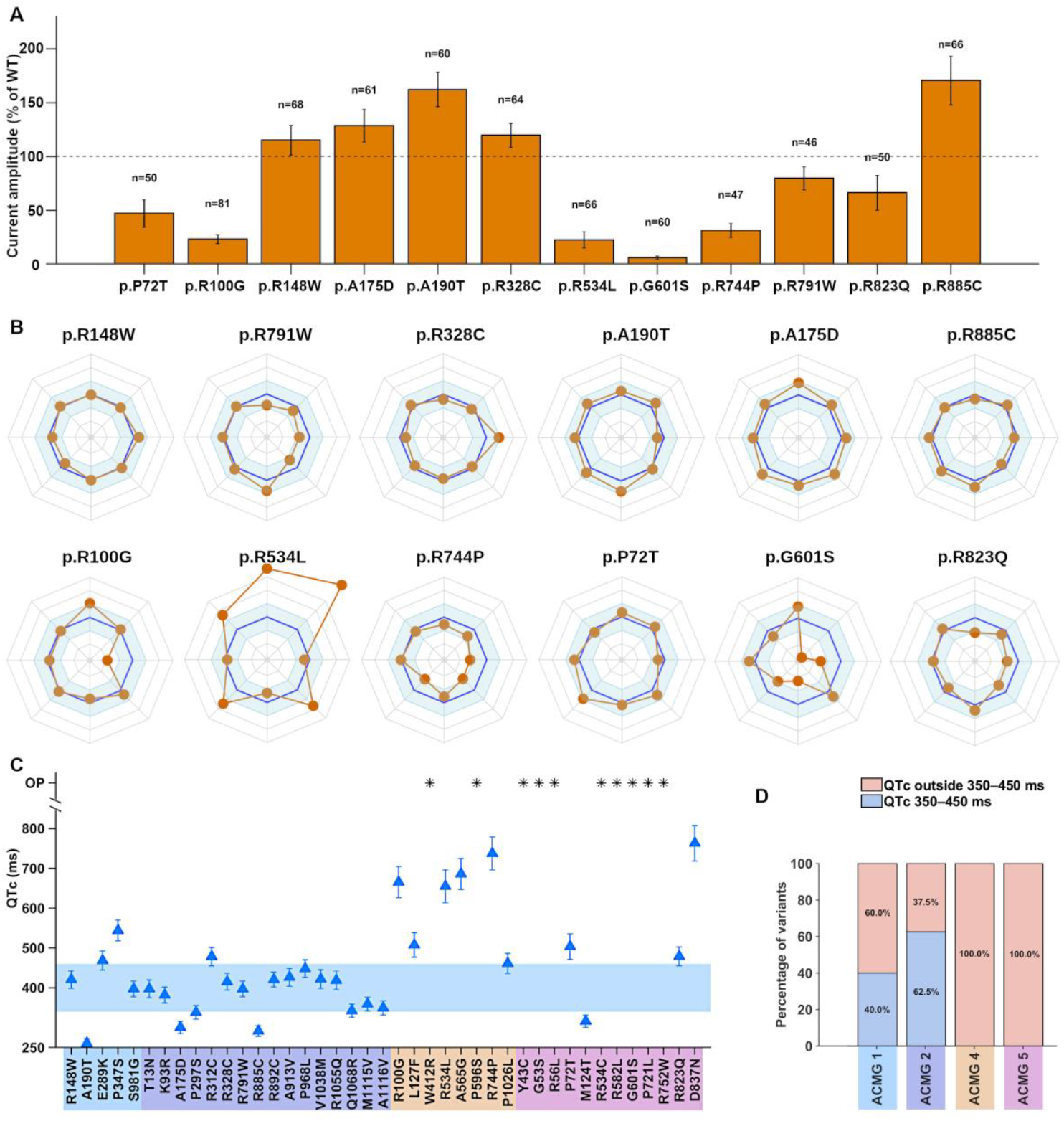
Functional screening framework for the determination of intrinsic pathogenicity of hERG variants. **A**, Bar plots illustrating current amplitudes of 12 representative hERG variants (6 from ACMG classes 1 and 2, 3 from ACMG class 4 and 3 from ACMG class 5), expressed as percentages of the WT. **B**, Radar plots of the 12 representative hERG variants shown in (**A**). These plots demonstrate the loss and gain of function mutations on the different hERG biophysical parameters, other than the current amplitude. **C**, QTc values derived from pseudo-ECGs generated by MeTAL for 41 hERG variants (including the ones shown in **A, B**) in the homozygous condition. Each filled blue triangle represents the mean QTc value (error bar shows the SD) obtained from 10 000 realizations across 1100 virtual patients. OP indicates *obviously pathogenic*, defined as pseudo-ECGs exhibiting abnormal features (ie, an abnormal number of ECG biomarkers compared with normal ECGs) and indicated by *. **D**, Distribution of predicted QTc values for the 41 hERG variants shown in (**C**), classified as within or outside the clinically normal QTc range (all sex confounded) and grouped according to ACMG pathogenicity class.

We then used MeTAL to generate virtual ECGs under homozygous conditions and compute the corresponding QTc values. While these simulated ECGs and QTc values do not have direct clinical relevance, since hERG variants are not expressed in a homozygous state in patients, this approach enables isolation of the intrinsic effects of the mutation. Specifically, it allows assessment of whether the mutation alone, in the absence of compensation by a wild-type allele, is sufficient to shift QTc beyond the clinical reference range.

Our findings showed that variants from classes 4-5 generally yielded pseudo-QTc values outside the physiological range, whereas variants from classes 1-2 showed more variable behavior (**Figure 6C, D**). A subset of variants also produced markedly abnormal ECG morphologies, including absent or duplicated biomarkers and fragmented or low-amplitude waveforms, precluding reliable QTc estimation. These variants were therefore designated as overtly pathogenic (OP) based on their severely abnormal simulated electrophysiological behavior.

Overall, this prescreening enabled the identification of 27 out of 41 variants for further investigation under heterozygous conditions and subsequent comparison with patient data.

### MeTAL recapitulates clinical QTc phenotypes across KCNH2 variants

To assess clinical relevance, we next performed high-throughput profiling of the 27 selected variants under heterozygous conditions (**Figure 7A-B, S7**). The resulting electrophysiological data (mean ± SEM for each biophysical parameter) were integrated into the MeTAL pipeline to simulate ECGs across a population of 11 million virtual patients, each harboring the same variant.

**Figure 7.**
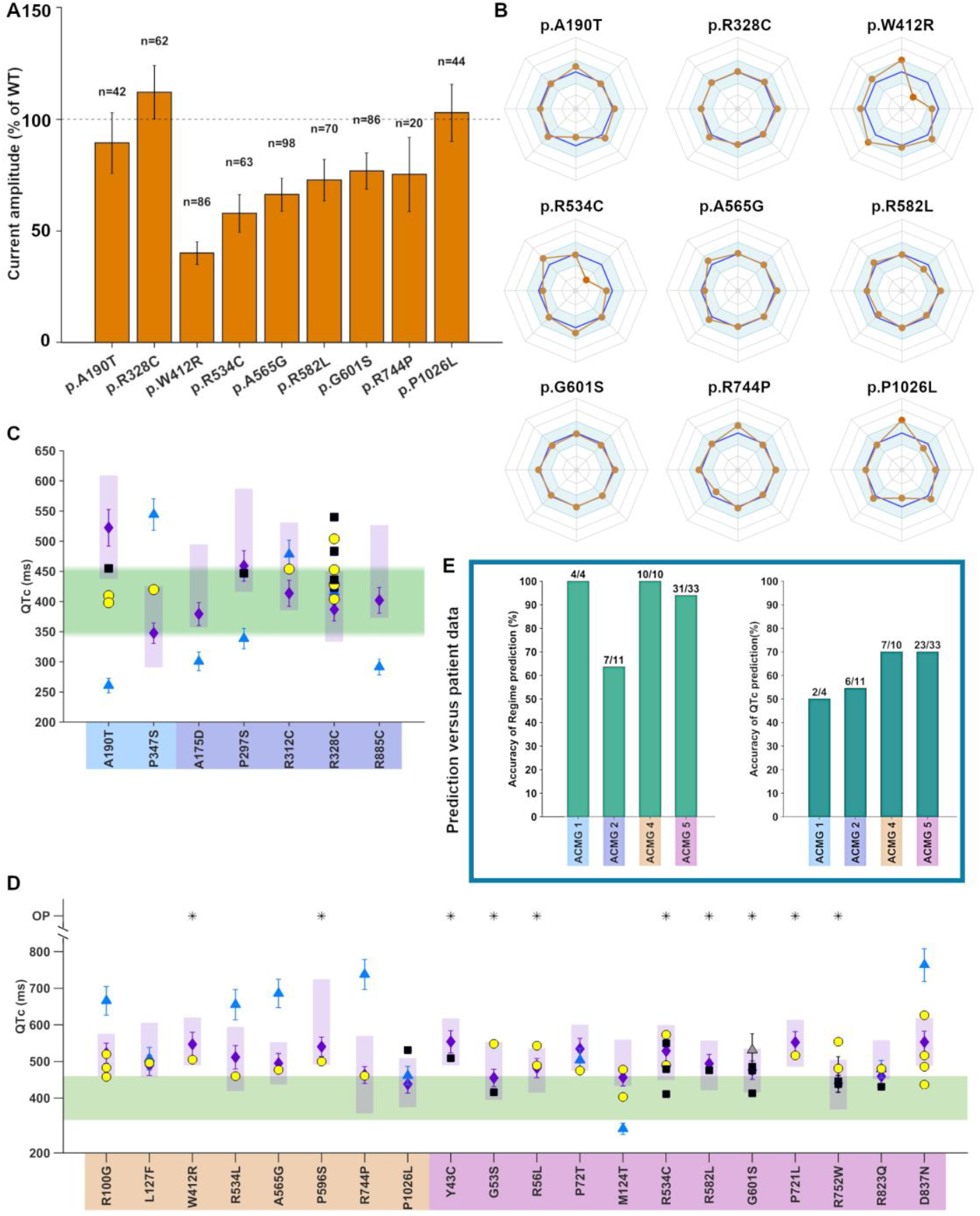
Functional screening–based prediction of intrinsic pathogenicity of hERG variants using the MeTAL algorithm. **A**, Bar plots illustrating current amplitudes, expressed as percentages of WT, for 9 representative variants selected from 27 prescreened hERG variants (ACMG classes 1–2, n=2; class 4, n=4; class 5, n=3). **B**, Radar plots of these variants showing loss- and gain-of-function effects across hERG biophysical parameters beyond current amplitude. **C–D**, QTc values derived from pseudo-ECGs generated by MeTAL for prescreened variants in the heterozygous condition, including 7 benign variants (**C**) and 20 pathogenic/likely pathogenic variants (**D**), overlaid with patient data. As in Figure 6C, blue triangles indicate simulated mean QTc values in the homozygous condition, and purple diamonds indicate simulated mean QTc values in the heterozygous condition. Yellow circles and black squares denote QTc values measured in female and male patients, respectively. Faded purple rectangles represent predicted QTc ranges in the heterozygous condition when experimental standard errors are incorporated into biophysical parameters. Overtly pathogenic (OP) variants, defined by pseudo-ECGs exhibiting abnormal features, are indicated by *. Simulated data are sex-confounded and grouped according to ACMG class. **E**, Statistical comparison of simulation-based predictions with clinical QTc data demonstrates high accuracy of MeTAL for intrinsic pathogenicity classification in ACMG classes 1, 4, and 5 under heterozygous conditions.

Simulated QTc distributions generally overlapped with observed patient data (**Figure 7C, D**). For instance, p.R752W showed a close agreement between simulated and clinical QTc values across a large cohort. In several additional cases—including p.P347S, p.R312C, p.R744P, and p.R823Q—variants that appeared markedly abnormal under homozygous conditions exhibited near-normal QTc behavior in heterozygous simulations, consistent with clinical observations.

Overall, MeTAL correctly captured the QTc regime (normal, prolonged, or shortened) across a substantial proportion of variants, with correct classification in 2/4 cases for class 1, 6/11 for class 2, 7/10 for class 4, and 23/33 for class 5 (**Figure 7E**). Simulated ECG waveforms were also consistent with patient recordings in both morphology and timing (**Figure 8A-B**), supporting the physiological fidelity of the framework.

**Figure 8.**
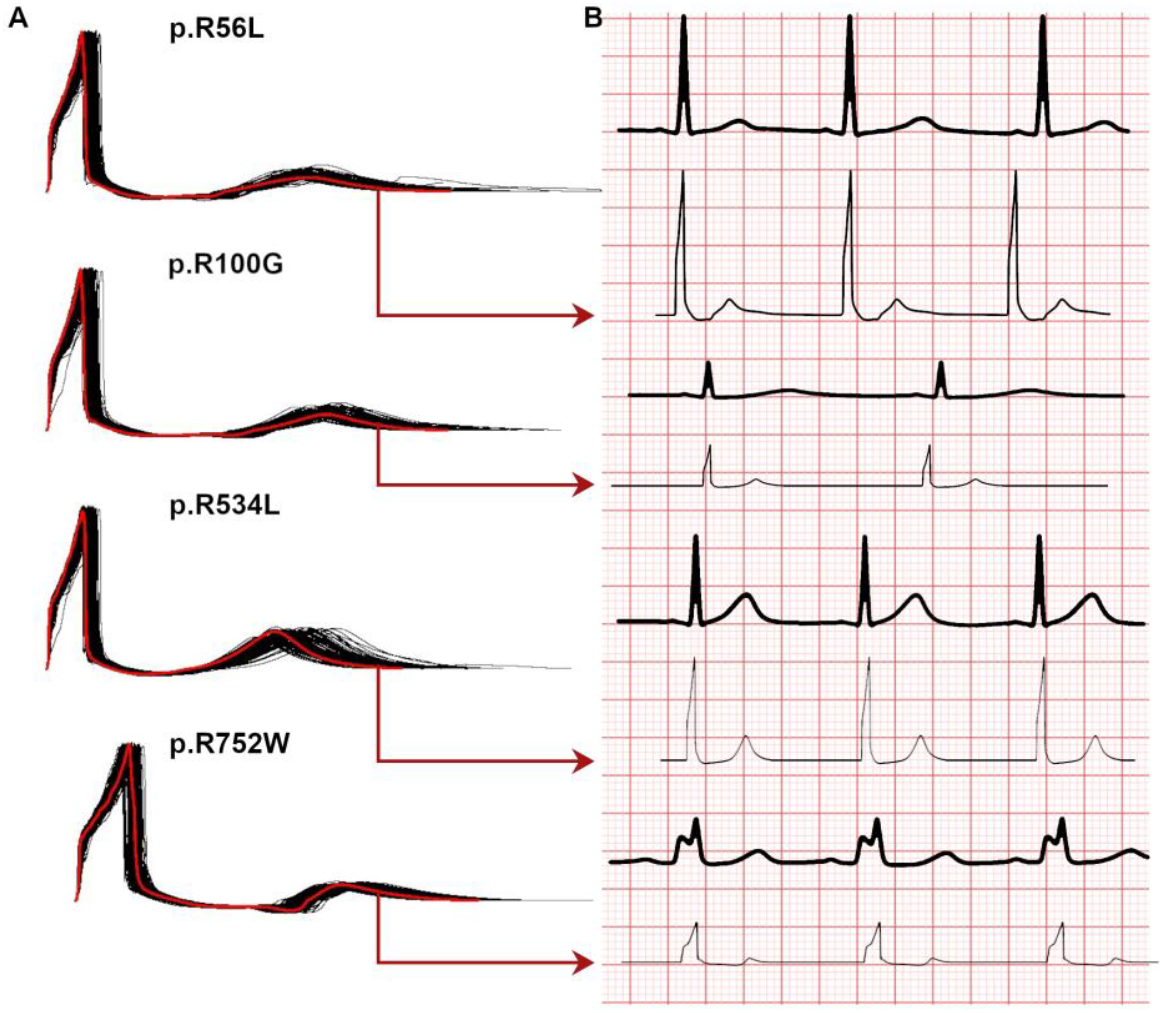
Agreement between simulated and patient ECG waveforms. **A**, Representative set of 500 pseudo-ECGs generated using hERG biophysical parameters corresponding to variants p.R56L, p.R100G, p.R534L, and p.R752W; the pseudo-ECG best matching the patient waveform is highlighted in red. **B**, Comparison between patient ECG recordings (bold line) and the red highlighted simulated ECG from A (now shown as thin black traces) at matched heart rate.

## Discussion

In this study, we developed a multiscale, biophysically detailed framework that links multiparametric hERG channel dysfunction to ventricular repolarization and ECG phenotypes. By integrating a flexible hERG formulation into a human ventricular electrophysiology model and combining it with a physiologically calibrated pseudo-ECG pipeline, MeTAL enables systematic evaluation of how distinct channel properties shape QT interval dynamics across scales.

A central finding of this work is that hERG variant effects are inherently multidimensional and cannot be adequately captured by conductance alone. Although reductions in maximal conductance (G_max_) are commonly used to assess *KCNH2* pathogenicity,^23,24^ our results demonstrate that other biophysical parameters – particularly those governing inactivation – play a dominant role in determining repolarization behavior. Both single-parameter and combinatorial analyses revealed that relatively modest perturbations in inactivation properties, including shifts in V_1/2,inact_ and changes in the slope of inactivation (*k*_inact_), can produce substantial alterations in QT duration, even in the presence of preserved or increased conductance. These findings establish a hierarchy of parameter importance in which inactivation dynamics emerge as key determinants of QT sensitivity.

Importantly, our results also show that interactions between parameters are highly nonlinear. The effect of G_max_ on QT duration depends strongly on the underlying configuration of gating parameters, such that similar levels of hERG current can give rise to markedly different repolarization phenotypes. This provides a mechanistic basis for the heterogeneous functional signatures observed across *KCNH2* variants and underscores the limitations of single-metric classification strategies.

By leveraging multiparametric electrophysiological data, MeTAL enables a direct mapping between experimentally measured channel properties and emergent ECG phenotypes. In a cohort of 41 *KCNH2* variants, simulated QTc distributions were consistent with observed patient values, and the model accurately captured the QTc regime (normal *versus* prolonged or shortened) across ACMG classes. Notably, several variants exhibiting pronounced dysfunction in homozygous simulations showed near-normal QTc behavior under heterozygous conditions, consistent with clinical observations. These results highlight the importance of considering allelic context and demonstrate that multiscale modeling can provide a physiologically grounded framework for interpreting variant effects on repolarization.

Automated patch-clamp (APC) assays, calibrated functional pipelines, and deep-mutational trafficking screens provide substantially richer functional datasets than conductance-based (G_max_) measurements alone.^2,13^ These approaches yield multiparametric electrophysiological profiles that are often incorporated into ACMG frameworks as functional evidence (PS3/BS3). Compared with single-metric assays, they offer higher throughput, improved reproducibility, and capture a broader spectrum of channel dysfunction. However, in many workflows, these multidimensional data are ultimately reduced to binary classifications (functionally normal *versus* abnormal) for clinical interpretation. As a result, mechanistic links between multiparametric channel dysfunction and downstream ECG phenotypes remain limited. In this context, MeTAL provides a complementary framework that integrates multiparametric electrophysiological inputs within a physiologically grounded multiscale model to generate QTc distributions and ECG phenotypes. This approach enables direct translation of experimental channel measurements into clinically interpretable repolarization outcomes.

Sequence- and ensemble-based pathogenicity predictors (e.g., REVEL, CADD, PolyPhen-2, CardioBoost, SPARC) take a fundamentally different approach, leveraging machine-learning or conservation- and structure-based features to generate variant pathogenicity scores. These methods provide rapid, genome-wide coverage and are widely used in diagnostic pipelines as supporting evidence, with disease-specific models such as CardioBoost improving performance for cardiac genes.^25,26^ However, they do not incorporate functional electrophysiology or ECG phenotypes, limiting mechanistic insight into variant effects. Moreover, performance on ion channel genes remains variable, with reported false positives and negatives for *KCNH2*.^26^ In contrast, MeTAL provides a biophysically grounded framework that links channel gating to electrophysiological phenotypes across biological scales, from cellular dynamics to tissue-level repolarization and ECG readouts. This approach enables identification of specific biophysical mechanisms underlying QTc alterations, complementing existing predictive tools with mechanistic interpretability.

The present study also has implications beyond variant interpretation. Because MeTAL explicitly captures the contribution of multiple channel properties, it provides a platform for evaluating how pharmacological modulation of hERG function may influence repolarization beyond simple current block, potentially refining drug safety assessment. Multiscale cardiac models are central to drug cardiotoxicity pipelines, where they are used to simulate action potentials and ECGs and to assess QT prolongation and arrhythmogenic risk. However, conventional approaches typically represent drug effects primarily through ionic current block, without accounting for potential modulation of other channel properties.^27^ Many existing models also rely on simplified channel formulations – such as instantaneous inactivation, single-conductance scaling, or Hodgkin-Huxley gating with limited flexibility – which are not designed to incorporate detailed multiparametric descriptions of channel behavior.^28^ MeTAL extends these approaches by integrating an augmented *I*_*Kr*_ formulation into the ORd model that decouples kinetics from voltage dependence and explicitly captures inactivation dynamics. In combination with a physiologically calibrated pseudo-ECG transformation, this framework enables incorporation of multiparametric electrophysiological data into multiscale simulations and facilitates evaluation of their impact on QT behavior.

By moving beyond conductance-based representations, this approach provides a foundation for assessing how partial or multidimensional modulation of hERG channel properties may influence repolarization, with potential implications for refining drug safety evaluation.

Several limitations should be acknowledged. The pseudo-ECG transformation relies on a segment-wise rescaling procedure calibrated to physiological reference ranges and assumes that similar transformations apply across normal and abnormal conditions. While this approach enables alignment of model outputs with clinically relevant ranges, it may not fully capture condition-specific variations in ECG morphology. In addition, automated extraction of ECG biomarkers depends on consistent feature identification and may differ from manual clinical annotation. Future development of standardized and automated ECG delineation methods, together with larger disease-specific datasets, could further improve the robustness and clinical applicability of the framework.

In conclusion, this work demonstrates that accurate interpretation of *KCNH2* variant effects requires a multiparametric and mechanistically grounded framework. By linking detailed channel kinetics to ECG phenotypes across scales, MeTAL provides a scalable approach for integrating experimental electrophysiology with clinically relevant outcomes. These findings support the systematic characterization of hERG gating properties beyond conductance and establish a foundation for improved interpretation of variant pathogenicity and proarrhythmic risk.

## Methods

### Clinical evaluation

This study agreed with the local guidelines for genetic research and has been approved by the local ethical committees. Informed, written consent was obtained from each patient who agreed to participate in the study. The Bamacoeur database from the French clinical network against inherited cardiac arrhythmias (Cardiogen) was prospected. Standard 12-lead electrocardiography recordings were conducted. Patients were considered affected when LQTS symptoms were associated with QTc exceeding 450 and 460 ms for men and women, respectively.

### Mutation analysis

Genomic DNA was extracted from peripheral blood lymphocytes using standard protocols. The patients were screened for mutations in, at least, the 3 major genes involved in the long QT syndrome, *KCNQ1, KCNH2*, and *SCN5A*.

### Cell Culture and Transfection

HEK293 cells were cultured as previously described^13^. The hERG channel (NCBI: NM_000238.4) was cloned into pCDNA5/FRT/TO and mutated using the Gibson assembly method. Plasmids were introduced into HEK293 cells by electroporation using the MaxCyte STx system (OC-100 cassettes, MaxCyte Inc., MD, USA)^19^, in homozygous (100%) and heterozygous conditions (50% wild-type and 50% variant). After 24 hrs, cells were trypsinized and resuspended in external NMDG solution (in mmol/L: 80 NaCl, 4 KCl, 2 CaCl_2_, 1 MgCl_2_, 5 glucose, 60 N-Methyl-D-Glucamine (NMDG), 10 HEPES - pH 7.4 with NaOH, 280 ± 3 mOsm) for hERG current recordings.

### Automated patch-clamp protocol

We adapted a manual patch-clamp voltage protocol for high-throughput current recording using the Nanion SyncroPatch 384PE^13^, incorporating six sub-protocols to record all biophysical parameters of hERG channel. Key modifications included increased external Ca^2+^ (2.48 mmol/L) and internal fluoride (110 mmol/L), requiring a +30-mV shift in prepulse potentials to compensate for the calcium-induced shift in activation and inactivation V_0.5_^29^. The holding potential was –80 mV, and the full recording protocol lasted under 90 s. Recordings used external NMDG solution and internal solution containing (in mmol/L) 110 KF, 10 NaCl, 10 KCl, 10 EGTA, and 10 HEPES (280 ± 3 mOsm, pH 7.2 with KOH; see Ref.^20^ for more details), at room temperature.

### Data analyses

Automated R scripts were used for analysis^13^. Quality control admitted cells with access resistance, Ra < 10 MΩ, seal resistance, Rs > 600 MΩ, leak current amplitude measured at -80 mV between 0 and -200 pA, and < 20% current run-down during each sub-protocol, the longest being less than 35 s. Except for G_max_, only 500 pA–10 nA currents were analyzed for the biophysical parameters, to avoid inaccurate analysis due to insufficient current or insufficient clamp^30^. Parameters were compared to wild-type controls recorded on the same day. Statistical tests included parametric tests and ANOVA. Significance: p < 0.05 (*), p < 0.01 (**), p < 0.001 (***). For further details on radar plot design, see **Supplementary Methods**.

### Mathematical model for human ventricular cardiomyocyte

The human ventricular cardiomyocyte model used in this study is based on the O’Hara–Rudy (ORd)^9^ formulation, with a modified representation of the rapid delayed rectifier potassium current (*I*_*Kr*_, hERG). Electrical activity of each cardiomyocyte was simulated by solving Eq.12.

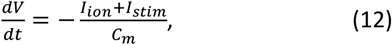

Here, *V* is the transmembrane potential, *I*_*stim*_, the external stimulus current, and *I*_*ion*_, the total ionic current produced by the cell, evaluated as a sum of 15 ionic currents (Eq.13).

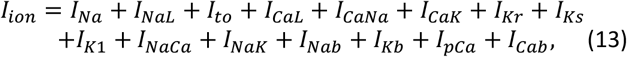

Full mathematical details of the hERG current formulation are provided in **Results** and **Supplementary Methods**.

### Pseudo-ECG generation

Pseudo-ECGs were generated using a one-dimensional transmural ventricular strand model adapted from the Gima-Rudy formulation.^16^ The strand comprised subendocardial, midmyocardial, and subepicardial cells. Each strand was electrically paced at 1 Hz from the sub-endocardial end. Electrical activity along the strand was computed by solving a reaction-diffusion equation (Eq.14):

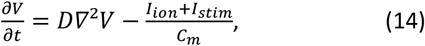

Where *D* controls the speed of propagation of the signal down the strand. The resulting extracellular unipolar potential (ϕ_*e*_) was computed at a hypothetical electrode positioned 2 cm away from the sub-epicardial end of the strand along the longitudinal axis, according to Eq.15.^16^

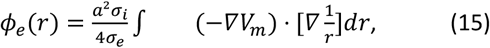

### Evaluation of single-parameter perturbations

To assess the electrophysiological impact of individual hERG parameters, pseudo-ECGs were simulated by varying one parameter at a time while keeping all other model parameters fixed in a one-dimensional ventricular strand configuration. Parameter perturbations were explored over predefined ranges for each of the nine hERG properties, including conductance scaling, voltage shifts, and kinetic ratios. Specifically, G_max_ was varied between 0 and 2, *k*^inact^ between 0.5 and 2.5, other kinetic parameters between 0.25 and 5, and voltage-dependent parameters within ±25 mV. This approach enabled systematic evaluation of the electrophysiological consequences of individual parameter variations across a broad range of physiologically plausible conditions.

## Acknowledgements

We acknowledge Dr Imani Younoussa (*l’institut du thorax*, Nantes) for his expert patients’ ECG analysis, Dr. Florence Kyndt (*l’institut du thorax*, Nantes) and Aurélie Loriou (Centre de référence des troubles du rythme cardiaque héréditaires ou rares de l’Ouest, CHU Nantes) for their digilent clinical data management.

## References

1. Vandenberg, J. I. et al. hERG K+ channels: structure, function, and clinical significance. Physiological reviews (2012).

2. Kozek, K. A. et al. High-throughput discovery of trafficking-deficient variants in the cardiac potassium channel Kv11.1. Heart rhythm 17, 2180–2189 (2020).

3. Ng, C.-A. et al. High-throughput phenotyping of heteromeric human ether-à-go-go-related gene potassium channel variants can discriminate pathogenic from rare benign variants. Heart rhythm 17, 492–500 (2020).

4. Thomson, K. L. et al. Clinical interpretation of KCNH2 variants using a robust PS3/BS3 functional patch-clamp assay. Human Genetics and Genomics Advances 5 (2024).

5. O’Neill MJ, et al. Multiplexed assays of variant effect and automated patch clamping improve KCNH2-LQTS variant classification and cardiac event risk stratification. Circulation 150:1869–1881 (2024).

6. Ma JG, Vandenberg JI, & Ng CA. Development of automated patch clamp assays to overcome the burden of variants of uncertain significance in inheritable arrhythmia syndromes. Front Physiol. 14:1294741 (2023).

7. Hodgkin AL, Huxley AF. A quantitative description of membrane current and its application to conduction and excitation in nerve. J Physiol. 117(4):500–44 (1952).

8. Lampert, A. & Korngreen, A. Markov modeling of ion channels: implications for understanding disease. Progress in molecular biology and translational science 123, 1–21 (2014).

9. O’Hara, T., Virág, L., Varró, A. & Rudy, Y. Simulation of the undiseased human cardiac ventricular action potential: model formulation and experimental validation. PLoS computational biology 7, e1002061 (2011).

10. Grandi, E., Pasqualini, F. S., Puglisi, J. L. & Bers, D. M. A novel computational model of the human ventricular action potential and Ca transient. Biophysical journal 96, 664a–665a (2009).

11. Ten Tusscher, K. H. & Panfilov, A. V. Alternans and spiral breakup in a human ventricular tissue model. American Journal of Physiology-Heart and Circulatory Physiology 291, H1088–H1100 (2006).

12. Alameh, M. et al. A need for exhaustive and standardized characterization of ion channels activity. The case of KV11. 1. Frontiers in Physiology 14, 1132533 (2023).

13. Oliveira-Mendes, B. et al. A standardised hERG phenotyping pipeline to evaluate KCNH2 genetic variant pathogenicity. Clinical and Translational Medicine 11, e609 (2021).

14. Jiang, C. et al. A calibrated functional patch-clamp assay to enhance clinical variant interpretation in KCNH2-related long QT syndrome. The American Journal of Human Genetics 109, 1199–1207 (2022).

15. Lei, C. L. et al. Rapid characterization of hERG channel kinetics I: using an automated high-throughput system. Biophysical Journal 117, 2438–2454 (2019).

16. Gima, K. & Rudy, Y. Ionic current basis of electrocardiographic waveforms: a model study. Circulation research 90, 889–896 (2002).

17. Moss, A. J. et al. Increased Risk of Arrhythmic Events in Long-QT Syndrome With Mutations in the Pore Region of the Human Ether-a-go-go–Related Gene Potassium Channel. Circulation 105(7), 794–799 (2002).

18. Gustina, A. S. & Trudeau, M. C. The eag domain regulates hERG channel inactivation gating via a direct interaction. Journal of General Physiology 141, 229–241 (2013).

19. Zhou, Z. et al. Properties of HERG channels stably expressed in HEK 293 cells studied at physiological temperature. Biophysical journal 74, 230–241 (1998).

20. Drouin, E., Charpentier, F., Gauthier, C., Laurent, K. & Le Marec, H. Electrophysiologic characteristics of cells spanning the left ventricular wall of human heart: evidence for presence of M cells. Journal of the American College of Cardiology 26, 185–192 (1995).

21. Jalife, J. & Stevenson, W. G. Zipes and Jalife’s Cardiac Electrophysiology: From Cell to Bedside, E-Book (Elsevier Health Sciences, 2021).

22. Normal ECG. e-cardiogram.com. https://www.e-cardiogram.com/ecg-normal/ (accessed July 21, 2025).

23. Jiang, C., Richardson, E., Farr, J., Hill, A.P., Ullah, R. et al. A calibrated functional patch-clamp assay to enhance clinical variant interpretation in KCNH2-related long QT syndrome. The American Journal of Human Genetics, 109 (7), 1199–1207 (2022).

24. Thomson, K.L., Jiang, C., Richardson, E., Westphal, D.S., Burkard, T., et al. Clinical interpretation of KCNH2 variants using a robust PS3/BS3 functional patch-clamp assay. HGG Adv. 2024 Apr 11;5(2):100270 (2024).

25. Ioannidis, N.M., Rothstein, J.H., Pejaver, V., Middha, S., McDonnell, S.K., et al. REVEL: An Ensemble Method for Predicting the Pathogenicity of Rare Missense Variants. Am J Hum Genet. 99(4):877–885 (2016).

26. Zhang, X., Walsh, R., Whiffin, N., Buchan, R., Midwinter, W., et al. Disease-specific variant pathogenicity prediction significantly improves variant interpretation in inherited cardiac conditions. Genet Med. 23(1):69–79 (2021).

27. Heitmann, S., Vandenberg, J.I., Hill, A.P. Assessing drug safety by identifying the axis of arrhythmia in cardiomyocyte electrophysiology. eLife 12:RP90027 (2023).

28. Meunier, C. & Segev, I. Playing the Devil’s advocate: is the Hodgkin–Huxley model useful? Trends Neurosci. 25(11), 558–563 (2002).

29. Ho, W.-K., Kim, I., Lee, C. O. & Earm, Y. E. Voltage-dependent blockade of HERG channels expressed in Xenopus oocytes by external Ca2+ and Mg2+. The Journal of Physiology 507, 631–638 (1998).

30. Montnach, J., Lorenzini, M., Lesage, A., Simon, I., Nicolas, S., et al. Computer modeling of whole-cell voltage-clamp analyses to delineate guidelines for good practice of manual and automated patch-clamp. Sci Rep 11, 3282 (2021).

